# Chloroplast-derived hydrogen peroxide coordinates photosynthesis with stomatal opening

**DOI:** 10.1101/2025.03.27.645763

**Authors:** Georgia Taylor, Julia Walter, Johannes Kromdijk

**Affiliations:** Department of Plant Sciences, University of Cambridge, Cambridge, Cambridgeshire, CB23EA, UK; Carl R Woese Institute for Genomic Biology, University of Illinois at Urbana-Champaign, IL61801, Illinois, USA

## Abstract

Stomatal pores on plant leaves control the entry point of carbon into the terrestrial biosphere and concomitant release of water vapour into the atmosphere. Stomatal aperture changes need to be coordinated with photosynthesis to balance carbon gain against water loss. Despite this importance, molecular signals underpinning this coordination, and the cell types from which such signals originate, remain poorly understood. Here we show that chloroplast-derived H_2_O_2_ coordinates stomatal movements with photosynthesis. Specifically, when H_2_O_2_ scavenging is compromised in the cytosolic *ascorbate peroxidase1* (*apx1*) mutant, stomatal opening is increased. This is rescued when photosynthetic H_2_O_2_ production derived from plastoquinol is reduced either genetically or pharmacologically. Furthermore, cell-specific complementation of APX1 in the *apx1* mutant demonstrated that H_2_O_2_ signals from both guard cells and mesophyll cells contribute significantly to stomatal movements. Altogether, our findings identify H_2_O_2_ as a key photosynthesis-derived signal underpinning effective coordination between stomatal conductance and photosynthesis, a key prerequisite for terrestrial plant life, and may inspire new strategies to enhance photosynthetic water use efficiency in crop species.

## Introduction

Stomata are microscopic pores in the leaf epidermis which govern the diffusional exchange of CO_2_ and water between terrestrial plants and the surrounding atmosphere. Highly specialised guard cells regulate stomatal aperture via turgor changes in response to various environmental and endogenous cues. Stomatal opening and photosynthesis are coordinated by molecular signalling pathways in response to changes in intercellular CO_2_ (*C*_i_) as well as *C*_i_-independent responses^1,2^. Both of these contribute equally^3^ to a gradual stomatal opening response to light in the photosynthetically active part of the solar spectrum^1,4^, also termed the stomatal red light response.

The molecular mechanisms underpinning the *C*_i_-independent stomatal red-light response are poorly understood. Stomatal opening is abolished by inhibition of photosystem II (PSII) photochemistry by the herbicide 3-(3,4-dichlorophenyl)-1, 1-dimethylurea (DCMU)^4,5^, which blocks the Q_B_ site on PSII, providing a link with photosynthesis. Suppression of stomatal red light responses in *Arabidopsis* ‘crumpled leaf’ mutants, which lack guard cell chloroplasts^6^ provides further evidence for control by guard-cell photosynthesis. In addition, recent work demonstrated that red light-dependent phosphorylation of Threonine-881 of guard cell plasma membrane (PM) H^+^-ATPases, which leads to stomatal opening, is dependent on guard cell photosynthesis^7,8^. While a link with photosynthesis is thus well-established, the molecular signals have remained elusive. Based on a synthesis of empirical observations Busch (2014) postulated that the redox state of the chloroplast plastoquinone/plastoquinol (PQ(H_2_)) pool, which reflects the balance between PQ reduction at PSII and oxidation at the cytochrome *b*_6_*f* complex, might be involved as an early signal. While this provides a first clue, further mechanistic evidence for such a link is still lacking.

More recently, guard cell accumulation of H_2_O_2_ was found to play an essential role in light-induced stomatal opening^9^ by promoting the nuclear localisation of KIN10, a catalytic subunit of the energy sensor Sucrose Non-Fermenting 1 related protein kinase 1 (SnRK1), leading to activation of starch degradation pathways. This guard cell specific accumulation of H_2_O_2_ appears to be conserved across a wide evolutionary range of terrestrial plants, however the origin of H_2_O_2_ accumulated in the guard cells and the impact on coordination between stomatal movements and photosynthesis still remain unexplored.

The thylakoid reactions are the primary source of reactive oxygen species (ROS) production in photosynthesising cells^10,11^. Within the chloroplast, H_2_O_2_ is produced both via the dismutation of superoxide (O_2_^-^) by thylakoid-bound and stromal superoxide dismutase (SOD) enzymes^12,13^ and via the spontaneous reaction between plastoquinol (PQH_2_) and O_2_^-^ ^14–17^ Click or tap here to enter text. Notably, H_2_O_2_ production within the chloroplast PQ/PQH_2_ pool displays a stochiometric relationship with light intensity^18,19^. Thylakoid-generated H_2_O_2_ can be detected outside of the chloroplast, even at low light intensity^15^, which along with its relatively long half-life^20^, capacity to travel across membranes^15,21^, and established role as a signalling molecule in a range of cellular processes^20,22,23^, makes H_2_O_2_ a strong candidate for communicating photosynthesis-derived information throughout the cell.

While guard cells are photosynthetically active, their capacity is thought to be limited^24^ and the bulk of leaf photosynthetic capacity resides in mesophyll cells. This implies that coordination between photosynthesis and stomatal conductance requires a mesophyll-derived signal. Indeed, stomatal red light responses in isolated epidermal peels are often suppressed compared to intact leaves^25–27^, but the molecular nature of such mesophyll-based signals is unclear. However, it is well-known that mesophyll cells accumulate chloroplast-derived H_2_O_2_ in response to light^28^ and this could be communicated to guard cells. Intercellular communication of H_2_O_2_ signals can occur both via plasmodesmata^29^, aquaporins^30^ or potentially even via the vapour phase^31^. In addition to this ‘passive’ transport, RESPIRATORY BURST OXIDASE HOMOLOGs (RBOHs) are apoplastic NADPH-oxidases well-known to facilitate the amplification and cell-cell communication of H­_2_O_2_ signals^32^. Indeed, light-induced stomatal opening is suppressed in *rbohd rbohf* double mutants^9,33^.

In the present work, we use a combination of genetic and pharmacological perturbations of photosynthetic H_2_O_2_ production in plants lacking ASCORBATE PEROXIDASE 1 (*apx1*) to demonstrate that H_2_O_2_ is the elusive signalling molecule which connects photosynthesis with *C*_i_-independent stomatal red-light responses, which has been subject to debate for decades (reviewed by Lawson *et al.,* 2014). Our results further show that H_2_O_2_ provides a functional link between PQ redox state changes and stomatal movements, which can be perturbed by manipulation of guard cell H_2_O_2_ scavenging capacity. Finally, we show that cell-specific complementation of H_2_O_2_ scavenging capacity in guard cells and mesophyll cells in *apx1* rescues elevated stomatal responses and guard cell H_2_O_2_ accumulation, consistent with intercellular H_2_O_2_ signalling as a mechanism to reconcile contributions from both mesophyll and guard cells to stomatal red light responses. Altogether, these findings provide new understanding of the pivotal role of H_2_O_2_ signals underpinning effective coordination between stomatal conductance and photosynthesis, a key prerequisite for terrestrial plant life.

## Results

### Enhanced guard cell H_2_O_2_ accumulation and stomatal opening in apx1 are mitigated by overexpression of PsbS

To investigate putative connections between PQ redox state, photosynthetic H_2_O_2_ production and stomatal movements, red light-induced stomatal opening was analysed in the *apx1* mutant, which is deficient in cytosolic H_2_O_2_ scavenging and exhibits enhanced cytosolic H_2_O_2_ accumulation^34^, as well as in the L17 accession, which carries a second copy of Photosystem II subunit S (*PsbS*)^35^, possibly leading to suppression of photosynthesis-derived H_2_O_2_ production, and the *apx1*L17 double mutant. Red light dose response curves of net CO_2_ assimilation (*A_net_*; Fig. 1A) and stomatal conductance (*g_s_*; Fig. 1B) were measured while keeping *C*_i_ constant (Supplementary Fig. 1A). *A­_net_* increased hyperbolically with red light intensity and was significantly elevated in *apx1* above 200 µmol m^-2^ s^-1^ red light (*P*<0.05; Fig. 1A, see Supplementary Table 3 for full Dunnett’s comparisons). As observed previously^1–3^, *g_s_* increased significantly with red light intensity (*P*<0.05, Fig. 1B) despite keeping *C*_i_ constant. In the H_2_O_2_ scavenging mutant *apx1*, *g_s_* responded more strongly to increasing red light intensity compared to the other genotypes and was significantly higher than Col-0 above 200 µmol m^-2^ s^-1^ red light (*P*<0.05; Fig. 1B, see Supplementary Table 4 for full Dunnett’s comparisons), which is consistent with the elevated *A*_net_ for this genotype (Fig. 1A). In line with previous observations on *PsbS* overexpression^36,37^, *g_s_* in L17 was slightly lower compared to the other genotypes (Fig. 1B), although the observed decrease was not significant. The dampening effect of *PsbS* overexpression on *g_s_* was more pronounced in the double mutant *apx1*L17, where *g_s_* values were significantly decreased relative to *apx1* and were statistically indistinguishable from Col-0 (*P*>0.05), showing that overexpression of *PsbS* fully negated the impact of *apx1* on *g*_s_. Total stomatal opening response to red light in all genotypes was estimated as Δ*g*_s_ (= *g*_s max_ – *g*_s int_), where *g*_s max_ represents maximal *g_s_* at 800 µmol m^-2^ s^-1^ red light, and *g*_s int_ represents *g*_s_ measured after 2 hours dark adaptation (Fig. 1C). Interestingly, Δ*g*_s_ was significantly elevated in *apx1* compared to wild-type (*P*=0.003, see Supplementary Table 5 for all Dunnett’s comparisons). Since *g*_s int_ did not vary significantly across genotypes (Fig. 1B, Supplementary Table 4), this confirms that the *apx1* mutation increases stomatal sensitivity to red light intensity. These genotypic effects on *g*_s_ were specifically associated with altered regulation of stomatal opening rather than anatomical differences, since stomatal complex length and stomatal density were similar between genotypes (Supplementary Fig. 2).

**Figure 1.**
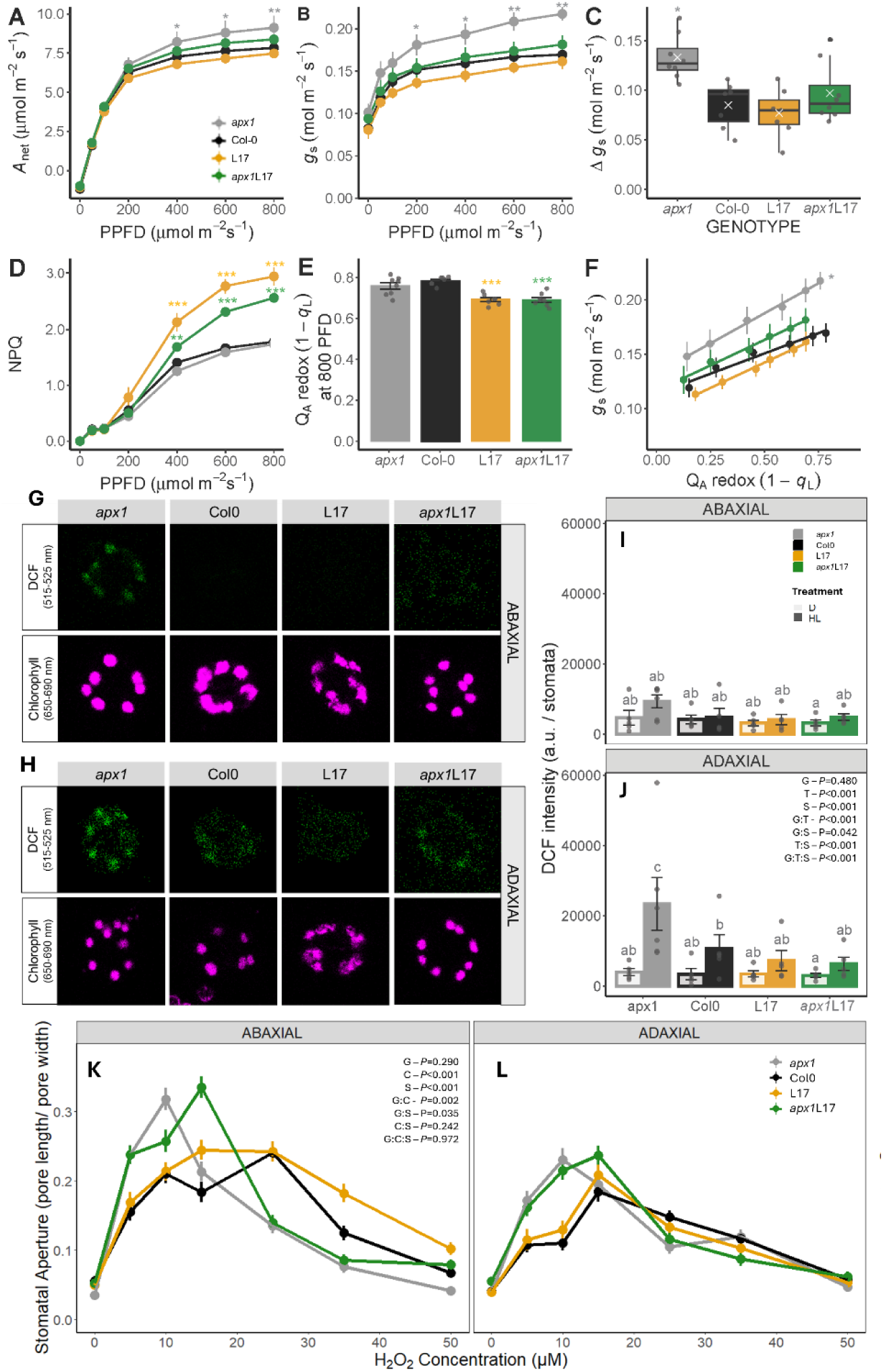
Rescue of elevated stomatal opening in *apx1* via PsbS overexpression. Leaf level gas exchange parameters (A) CO_2_ assimilation (*A*_net_) and (B) stomatal conductance (*g_s_*) were measured at a controlled intercellular CO_2_ (*C_i_*) concentration (∼300 µmol mol^-1^). (C) The total stomatal opening response across the entire red light response curve, estimated using Δ*g*_s_ *= g*_s max_ – *g*_s int_; where Δ*g*_s_ is the difference in stomatal conductance, *g*_s max_ is the maximum stomatal conductance measured at high red light (800 μmol m^-2^ s^-1^) and *g*_s int_ is the initial conductance measured after dark adaptation. Chlorophyll fluorescence was concomitantly measured after stabilisation at each light level to assess the effects of PsbS and APX1 levels on the response of the photosynthetic electron transport chain: (D) non-photochemical quenching (NPQ) and (E) the redox state of Quinone A (Q_A_), as estimated by the parameter 1–*q*_L_, at the end of the red light response curve (800 μmol m^-2^ s^-1^). (F) Linear regression analysis of the relationship between *g*_s_ and *Q*_A_ redox state to increasing red light under constant *C*_i._ Two-way repeated measures ANOVA tested the effects of genotype, red light intensity (PFD) and their interaction on the response curves. Asterisks represent significant differences compared to Col-0 (P<0.05) using a Dunnett’s pairwise comparison; significance codes, *P*<0.001 (***), <0.01 (**), <0.05 (*), n=7. The responses of DCF and chlorophyll fluorescence intensity were imaged via CLSM for (G) abaxial and (H) adaxial guard cells. DCF intensity was quantified in response to a 1 hour red light treatment (HL, 800 µmol m^-2^ s^-1^) or dark control for both abaxial (I) and adaxial (J) guard cells. n=5 biological replicates, >40 stomata measured per biological replicate. To establish the effects of PsbS and APX1 levels on the stomatal response to H_2_O_2_, isolated epidermal peels were dark adapted for 2 h before incubation with defined H_2_O_2_ concentrations for 2.5 hours in the absence of light. n=5 biological replicates, >40 stomata measured for each biological replicate for each leaf side. Three-way ANOVA tested the effects of genotype (G), leaf side (S), light treatment (T), and their interaction on the response of DCF, different letters represent significant differences between genotypes based on a Tukey’s comparison across all treatments and leaf sides (H & J). *N.B. For DCF, statistical analysis was performed on log transformed data*. Stomatal aperture (pore length/ pore width) was quantified for (K) abaxial and (L) adaxial surfaces in *Arabidopsis* single and double mutants. n=5 biological replicates, for each biological replicate >40 stomata were measured for each leaf side at each concentration. Error bars= ± SEM for all panels. Three-way ANOVA tested the effects of genotype (G), leaf side (S), H_2_O_2_ concentration (C), and their interaction on the response of stomatal aperture.

Next, we explored if the opposing effects of *apx1* and L17 on *g*_s_ responses to red light could be explained by contrasting effects on H_2_O_2_ accumulation. As expected, non-photochemical quenching (NPQ) was significantly increased in L17 and *apx1*L17 relative to Col-0 at high red light (>400 µmol m^-2^ s^-1^; Fig. 1D, see Supplementary Table 6 for Dunnett’s comparisons) and consequentially, the redox state of quinone A (Q_A_), as estimated by the chlorophyll fluorescence parameter 1 – *q*_L_, was significantly more oxidised in L17 and *apx1*L17 at 800 µmol m^-2^ s^-1^ red light (Fig. 1E, see Supplementary Table 7 for Dunnett’s comparisons). Consistent with previous observations^36^, a linear relationship between Q_A_ redox state and *g_s_* was observed across all genotypes (Fig. 1F). Multiple linear regression in combination with backwards stepwise elimination yielded a minimal model for the prediction of *g_s_ ,* which was able to capture a substantial proportion of the experimental variance (*F*(7, 172)=19.49, *P* <0.001, *R*^2^= 0.442, see Supplementary Table 8 for full regression analysis), based on significant additive effects of genotype and Q_A_ redox state. Notably, the y-intercept was significantly greater in the *apx1* H_2_O_2_ scavenging mutant than all other genotypes (*P*=0.048), such that *g*_s_ was consistently increased in *apx1* for any given PQ redox value, suggesting an increased sensitivity to red light intensity, likely driven by elevated cytosolic H_2_O_2_ levels.

Subsequently, we investigated whether these empirical relationships between Q_A_ redox state and *g_s_* could be explained by a mechanistic link between PQ redox state and guard cell H_2_O_2_ accumulation. Guard cell H_2_O_2_ accumulation, as estimated by dichlorofluorescein (DCF) intensity per stomatal complex, increased in response to the red light treatment relative to the dark control (Fig. 1G-I). In the illuminated samples, the DCF signal co-localised with chlorophyll autofluorescence, suggesting that the major fraction of light-driven H_2_O_2_ production originated from the chloroplasts. The extent of light-induced H_2_O_2_ accumulation was significantly higher in *apx1* (*P*<0.05, Fig 1H-J, see Supplementary Table 9 for Tukey’s comparisons), most clearly on the adaxial surface, likely due to the direction of illumination. Consistent with the opposing effects of *apx1* and L17 on *g_s_*, the increased guard cell H_2_O_2_ levels in *apx1* were also mitigated to wild-type levels in *apx1*L17. This shows that dampening the production of H_2_O_2_ from thylakoid reactions via *PsbS* overexpression was able to reverse the elevation in guard cell cytosolic H_2_O_2_ levels associated with the *apx1* mutation. The observed differences in DCF intensity could not be explained by differences in chlorophyll content, chloroplast area or positioning, which were similar between genotypes (Supplementary Fig. 3). To find out if stomatal sensitivity to H_2_O_2_ was affected by either mutation, stomatal aperture dose-responses to exogenous H_2_O_2_ were measured on isolated epidermal peels (Fig. 1K-L). Maximal stomatal aperture was approximately 20% higher in the abaxial surface than the adaxial surface across all genotypes (Fig. 1K-L). As expected, stomatal aperture responses to H_2_O_2_ were significantly influenced by *apx1,* as exemplified by the significant interaction between genotype and concentration (*F*(18,149)=2.46, *P*=0.002). Single and double mutants (*apx1* and *apx1*L17) were more sensitive to H_2_O_2_ concentration than genotypes carrying the wildtype *APX1* allele (Col-0 and L17), see Supplemental Table 10 for Tukey’s comparisons. This shows that reduced H_2_O_2_ scavenging capacity in *apx1* increases sensitivity of stomatal opening to H_2_O_2_ concentration in the buffer, whereas overexpression of *PsbS* associated with L17 had no impact on stomatal sensitivity to H_2_O_2_ and instead specifically affected production of thylakoid-derived H_2_O_2_.

### Multiple distinct sources of H_2_O_2_ are required for stomatal opening in response to red light

The impact of PsbS overexpression shows the importance of photosynthetically derived H_2_O_2_ in driving stomatal opening in response to red light. However, apoplastic H_2_O_2_ production, mediated by plasma membrane-bound NADPH oxidases such as RBOHD and RBOHF, also contributes to guard cell H_2_O_2_ accumulation and associated changes in stomatal aperture^38^. To disentangle the contributions of these H_2_O_2_ sources, we measured guard cell H_2_O_2_ accumulation and stomatal aperture in response to high-intensity red light (HL; 800 µmol m⁻² s⁻¹) following the infiltration of leaves with specific inhibitors targeting distinct H_2_O_2_ sources (Fig. 2). Mock-infiltrated samples exposed to HL showed significant elevation of H_2_O_2_ production estimated from DCF fluorescence, relative to samples kept in darkness (Fig. 2A-B). H_2_O_2_ production was completely suppressed in samples infiltrated with the herbicide 3-(3,4-dichorophenyl)-1,1-dimethylurea (DCMU), which maintains the chloroplast plastoquinone pool in a fully oxidised state, whereas infiltration with dibromothymoquinone (DBMIB), which maximally reduces the plastoquinone pool under light conditions, promoted H_2_O_2_ generation to levels slightly above mock-infiltrated samples. Infiltration with potassium iodide (KI) and N,N’-dimethylthourea (DMTU), generalist H_2_O_2_ scavengers, abolished H_2_O_2_ accumulation above dark control levels and so did infiltration with diphenyleneiodonium chloride (DPI) and imidazole, which inhibit apoplastic H_2_O_2_ formation by flavoproteins, such as RBOHs (Fig. 2A-D, see Supplementary Table 11 for Tukey’s comparison). Thus, inhibition of either apoplastic or photosynthetic H_2_O_2_ production prevents the increases in guard cell H_2_O_2_ levels in response to HL exposure.

**Figure 2.**
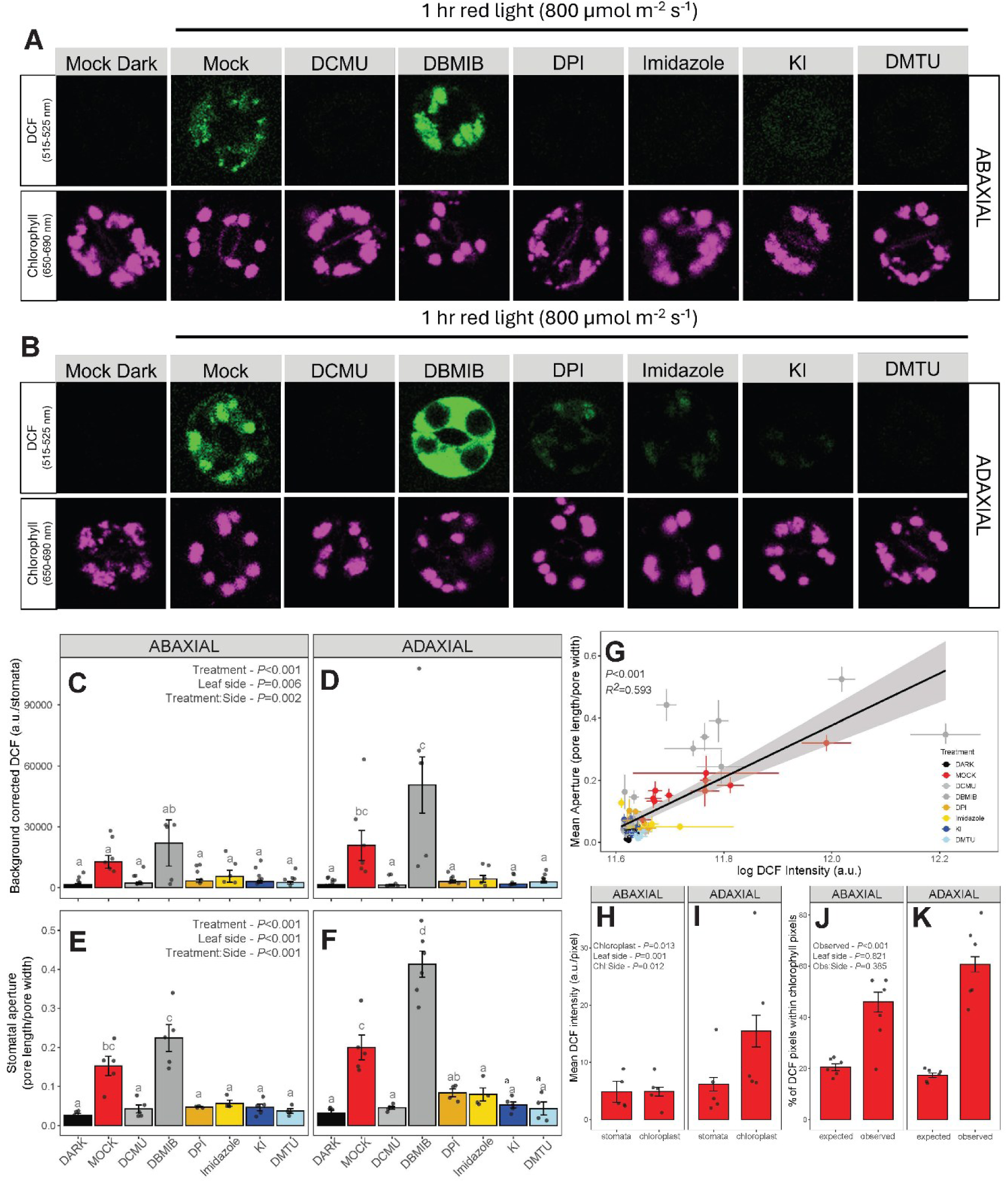
Manipulation of H_2_O_2_ accumulation modulates stomatal opening in response to red light. The effect of H_2_O_2_ inhibition from multiple sources on the response of stomatal aperture and H_2_O_2_ accumulation to high red light treatment (HL; 800 µmol m^-2^ s^-1^) in *Arabidopsis* whole leaves was investigated by imaging DCF and chlorophyll fluorescence in adaxial (A) and abaxial (B) guard cells via confocal microscopy. Guard cell DCF intensity was quantified in response to red HL in the presence of different inhibitors for both abaxial (C) and adaxial (D) guard cells. To assess the effect of H_2_O_2_ manipulation on stomatal opening, stomatal aperture (pore length/pore width) responses to red HL was quantified in the presence of different inhibitors for abaxial (E) and adaxial (F) guard cells. To investigate the coordination between stomatal opening and guard cell H_2_O_2_ levels a linear regression was performed between guard cell DCF intensity and stomatal aperture (G). Two-way ANOVA tested the effects of treatment and leaf side on the responses of DCF and stomatal aperture. Different letters represent statistically different Tukey’s pair wise comparisons (*P*<0.05). To further investigate the origin of guard cell H_2_O_2_, DCF intensity (a.u./ pixel) was analysed both at the whole stomata level and within only the chloroplast containing regions for each image for both abaxial (H) and adaxial (I) guard cells. Two-way ANOVA was used to test the effects of measurement region (chloroplast), leaf side, and their interaction on DCF intensity. Different letters represent statistically different Tukey’s pair wise comparisons (*P*<0.05). Additionally, the Mander’s coefficient was calculated to quantify the proportion of DCF pixels within chlorophyll containing pixels (%) for abaxial (J) and adaxial (K) guard cells, the expected % overlap is equal to the chlorophyll coverage (%) for each stoma, while the observed is the measured correlation coefficient for each stoma. Two-way ANOVA was used to measure the effect of observed vs expected, leaf side and their interaction on the degree of pixel overlap. (n=6 biological replicates, >40 stomata measured per replicate, error bars = SEM). *N.B. statistics for DCF intensity was performed on log transformed data*.

In line with significant H_2_O_2_ accumulation in mock-infiltrated samples, both abaxial and adaxial epidermal peels displayed significant stomatal opening in response to red light compared to dark conditions (Fig. 2E-F). Stomatal opening corresponded to a marked accumulation of H_2_O_2_ specifically within guard cells (Fig. 2A-D). Consistent with previous results (Fig. 1J), adaxial guard cells, which directly perceive the red light stimulus, exhibited higher DCF fluorescence than abaxial guard cells (Figs. 2A-D), and this was associated with greater stomatal apertures on the adaxial surface (Fig. 2E-F). Notably, red light-induced stomatal opening closely mirrored guard cell H_2_O_2_ accumulation patterns (Figs. 2C-D vs 2E-F). For all chemical treatments except DBMIB, the suppression of red light-induced H_2_O_2_ accumulation in guard cells blocked stomatal opening (Fig. 2E-F). Conversely, H_2_O_2_ accumulation (Fig. 2A-D) and stomatal opening (Fig. 2E-F) were both slightly elevated above wild-type levels in DBMIB-treated samples. Interestingly, all measurements of guard cell DCF accumulation and stomatal aperture could be described by a single highly significant positive correlation (F(1, 65)=116.5, *P*<0.001, *R*^2^=0.64; Fig. 2G see Supplementary Table 12 for full regression analysis), strongly supporting a relationship between guard cell H_2_O_2_ levels and stomatal opening.

To further verify the specific contribution of guard cell chloroplasts to guard cell cytosolic H_2_O_2_ levels, spatial variation in DCF intensity was assessed by comparing the DCF intensity averaged across the entire stomatal complex with the average from chlorophyll-containing regions, based on overlap with chlorophyll autofluorescence (Fig. 2H-I). In the absence of a significant contribution of guard cell chloroplasts to guard cell H_2_O_2_ levels, DCF fluorescence would be expected to show a uniform distribution across each stomatal complex. However, two-way ANOVA demonstrated significant effects of chloroplast-containing pixels (*F*(1, 16)=6.439, *P*=0.013), leaf side (*F*(1,16)=10.886, *P*=0.001), and their interaction on DCF intensity (*F*(1,16)=6.661, *P*=0.012). In agreement with the adaxial direction of illumination, DCF fluorescence was significantly elevated in chlorophyll-containing regions compared to whole stomata in adaxial guard cells (Fig. 2I, see Supplementary Table 13 for full Tukey’s comparisons). Consistently, spatial colocalization analysis of chlorophyll and DCF fluorescence channels estimated by the Mander’s coefficient^39^ demonstrated significantly higher overlap between both signals compared to the expected overlap with a spatially uniform DCF fluorescence signal (Figs. 2J-K; two-way ANOVA *F*(1, 16)=40.026, *P*<0.001), confirming that guard cell H_2_O_2_ predominantly originates from guard cell chloroplasts.

### Guard cell and mesophyll-targeted complementation of APX1 both significantly decrease red light-driven stomatal opening in apx1

The identification of H_2_O_2_ as a messenger molecule in stomatal red light responses provided an opportunity to investigate the specific cell types involved in coordinating this response, which is a long-standing question in stomatal biology (see review by Lawson *et al.,* 2014). *APX1* expression constructs were generated using cell-specific promoters and transformed into the *apx1* genetic background to modulate H_2_O_2_ scavenging capacity in guard cell-(pGC1^40^ and pMYB60^41^) or mesophyll-specific (pCytFBPase^42,43^) manner. All three cell-specific promoters were confirmed using promoter-eGFP fusions, showing strong GFP fluorescence from the targeted cell type, without detectable signal from the alternative cell type (Fig. 3A; Supplementary Fig. 4). Ubiquitous (pUBI::APX1) and mesophyll-specific (pCytFBPase::APX1) complementation led to strong accumulation of APX1 transcripts (Fig. 3B) and protein (Fig. 3C) in whole leaf extracts, which was much weaker in the guard-cell specific lines (pGC1::APX1 and pMYB60::APX1) due to the small contribution of guard cells to whole leaf extract. To assess the effect of cell-specific modulation of *APX1* expression on stomatal red light responses, red light response curves of *g*_s_ (Fig. 3D) and *A*_net_ (Fig. 3D) were measured at constant *C*_i_. The response of *g*_s_ varied significantly between genotypes (Fig. 3D). As observed previously (Fig. 1B), red light-induced stomatal opening was significantly elevated in *apx1* at high red light (Fig. 3D, *P*<0.05 Tukey’s). Notably, at light intensities >200 µmol m^-2^ s^-1^, *g*_s_ was significantly lower than *apx1* and Col-0 in all ubiquitous or guard cell-specific *APX1* complementation lines. Although less pronounced, *g*_s_ was also significantly lower in mesophyll-specific *apx1/*p*CytFBPase::APX1* lines relative to *apx1,* such that the response was similar to Col-0 across the entire red light response curve (Fig. 3D, see Supplementary Table 14 for Tukey’s comparisons). These results indicate that the H_2_O_2_-scavenging capacity of guard cells strongly affects the *C*_i_-independent red light response, and that H_2_O_2_ produced in mesophyll cells also contributes to stomatal opening. Interestingly, the strong decreases in *g_s_* due to ubiquitous and guard cell-specific APX1 complementation did not seem to affect *A*_net_ under constant *C*_i_, which remained similar between genotypes (Fig. 3E; *P*>0.05, Supplementary Table 15). Stomatal anatomy and density remained similar between genotypes (Supplemental Fig. 4), demonstrating that the observed changes in *g*_s_ resulted from altered stomatal regulation due to differences in cell-specific H_2_O_2_ scavenging capacity.

**Figure 3.**
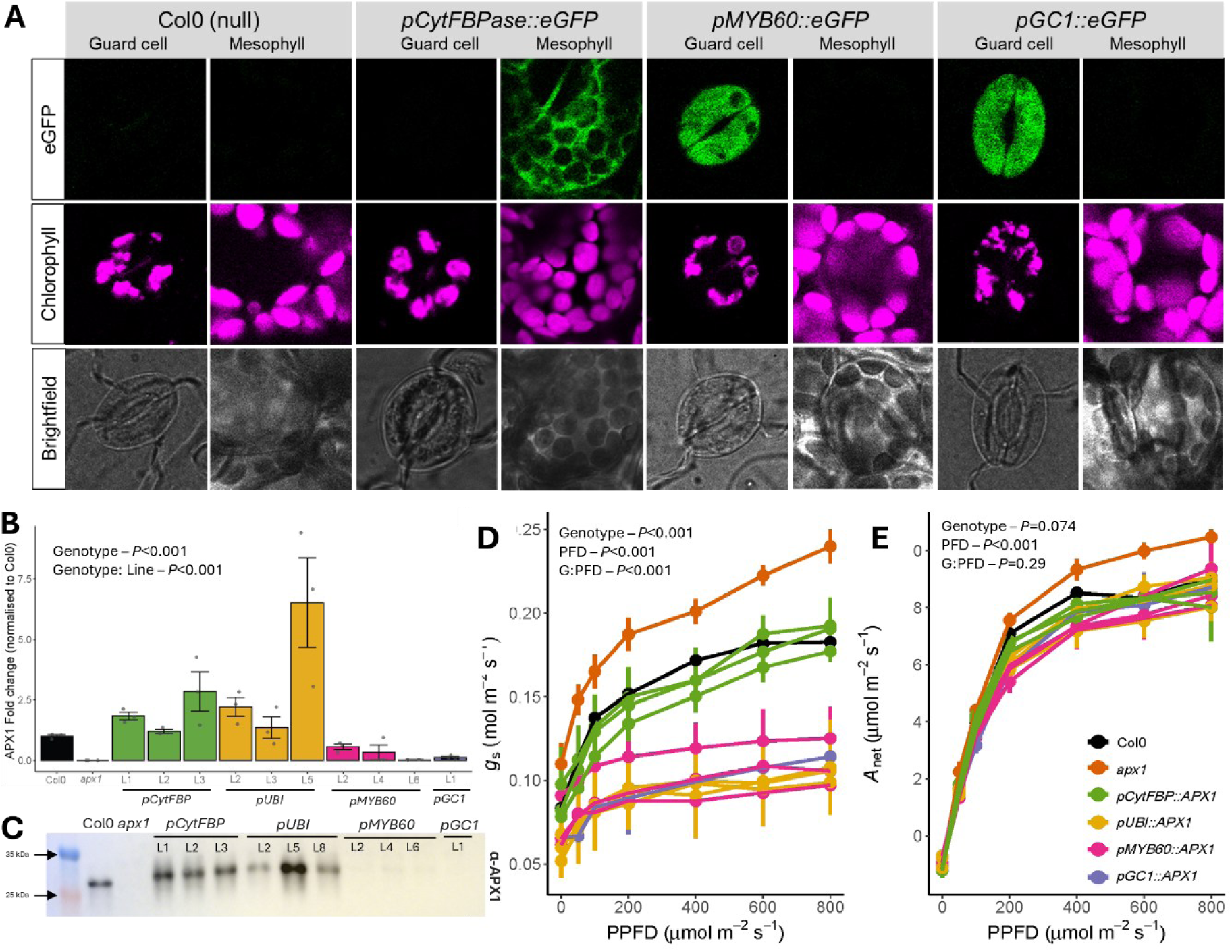
Rescue of elevated stomatal opening in *apx1* in cell-specific APX1-complemented lines. Cell specificity of APX1-complementation was verified. (A) CLSM imaging of epidermal and mesophyll layers in eGFP promoter-reporter lines confirmed mesophyll specific expression of the *pCytFBPase* and guard cell-specific expression of the *pGC1* and *pMYB60*. eGFP and chlorophyll autofluorescence were excited using a 488 nm laser and detected at 507 ±5 nm and 660 ± 25 nm, respectively. Expression patterns were verified for 5 independent insertion lines, representative images are shown. (B) RT-qPCR was used to verify *APX1* expression via mesophyll-specific (*pCytFBPase*) and ubiquitous (*pUBI)* complementation. Relative fold change was calculated using ΔΔCt and normalised against Col-0 (WT). *APX1* expression was quantified for three independent insertion lines for each promoter., n=4, error bars represent SEM. One way ANOVA tested the effect of genotype, with nested effects of line within genotype, on the expression of *APX1*. (C) Immunoblot against α-APX1 confirms the absence of APX1 in the *apx1* null, which is restored to at least wild-type levels via mesophyll or ubiqutious APX1-complementation. Leaf level gas exchange was used investigate the effect of cell-specific APX1-complementation on stomatal behaviour. The responses of (D) CO_2_ assimilation (*A*_net_) and (E) stomatal conductance (*g*_s_) were measured in response to increasing red light intensity at constant a intercellular CO_2_ concentration of ∼300 umol mol^-1^. Three independent insertion lines were measured for each promoter. n=7, error bars = SEM. Two-way repeated measures ANOVA tested the effects of genotype, red light intensity (PFD), and their interaction on red light response curves, with nested effects of line within genotype.

### Cell-specific APX1 complementation differentially impacts guard cell H_2_O_2_ accumulation and sensitivity

The observed suppression of stomatal opening via cell-specific complementation of *APX1* (Fig 3E) suggests that the stomatal response to red light is coordinated by guard cell H_2_O_2_ levels, which integrate H_2_O_2_ produced at local (e.g. guard cell chloroplasts) and distal sources. To empirically confirm this interpretation, the effects of cell-specific *APX1*-complementation on H_2_O_2_ accumulation were investigated. To investigate the impact of cell-specific APX1 complementation on mesophyll H_2_O_2_ levels, H_2_O_2_ was quantified using a plate-based Amplex Red assay on whole leaf discs (Supplemental Fig. 5). Under red light, Amplex Red fluorescence (background-corrected fluorescence units) was statistically similar to Col-0 in *pCytFBPase::APX1* and *pUBI::APX1* plants (Supplemental Fig. 5; *P*<0.05, Dunnett’s comparison), whereas H_2_O_2_ accumulation was significantly elevated in *apx1, pGC1::APX1* and *pMYB60::APX1* genotypes (Supplemental Fig. 5; *P*<0.05, Dunnett’s comparison), both cases are consistent with the cell-specificity of APX1 complementation. H_2_O_2_ accumulation in guard cells was analysed using DCF fluorescence intensity (Fig 4A-D). DCF intensity per stomatal complex increased in response to the red light illumination, with the DCF signal largely co-localising with chlorophyll autofluorescence (Fig. 4A-B). Consistent with previous observations (Fig. 1G-J), red light-induced H_2_O_2_ accumulation in both abaxial and adaxial guard cells was significantly elevated in *apx1* compared to Col-0 and all guard cell APX1-complementation genotypes (*apx1*/*pGC1*/*pMYB60*/*pUBI*::APX1; *P*<0.05, see Supplementary Table 16 for Tukey’s comparisons), demonstrating that guard cell-specific or ubiquitous complementation of APX1 successfully restored cytosolic H_2_O_2_ scavenging capacity in guard cells. An intermediate DCF intensity was observed for mesophyll-specific *apx1*/*pCytFBPase::APX1* plants, which was statistically similar to both the *apx1* parent genotype and Col-0 (Fig. 4C-D), in line with a contribution of mesophyll-derived H_2_O_2_ to guard cell H_2_O_2_ levels. Despite the substantial reduction in *g*_s_ compared to Col-0 (Fig. 3D), none of the complementation lines expressing *APX1* in guard cells (*apx1/pGC1/pMYB60/pUBI::APX1;* Fig. 4C-D) deviated significantly from Col-0 in guard cell H_2_O_2_ accumulation, which may be due to limited resolution but could also suggest that altered *g*_s_ responses in these plants also reflect altered H_2_O_2_ sensitivity of guard cells.

**Figure 4.**
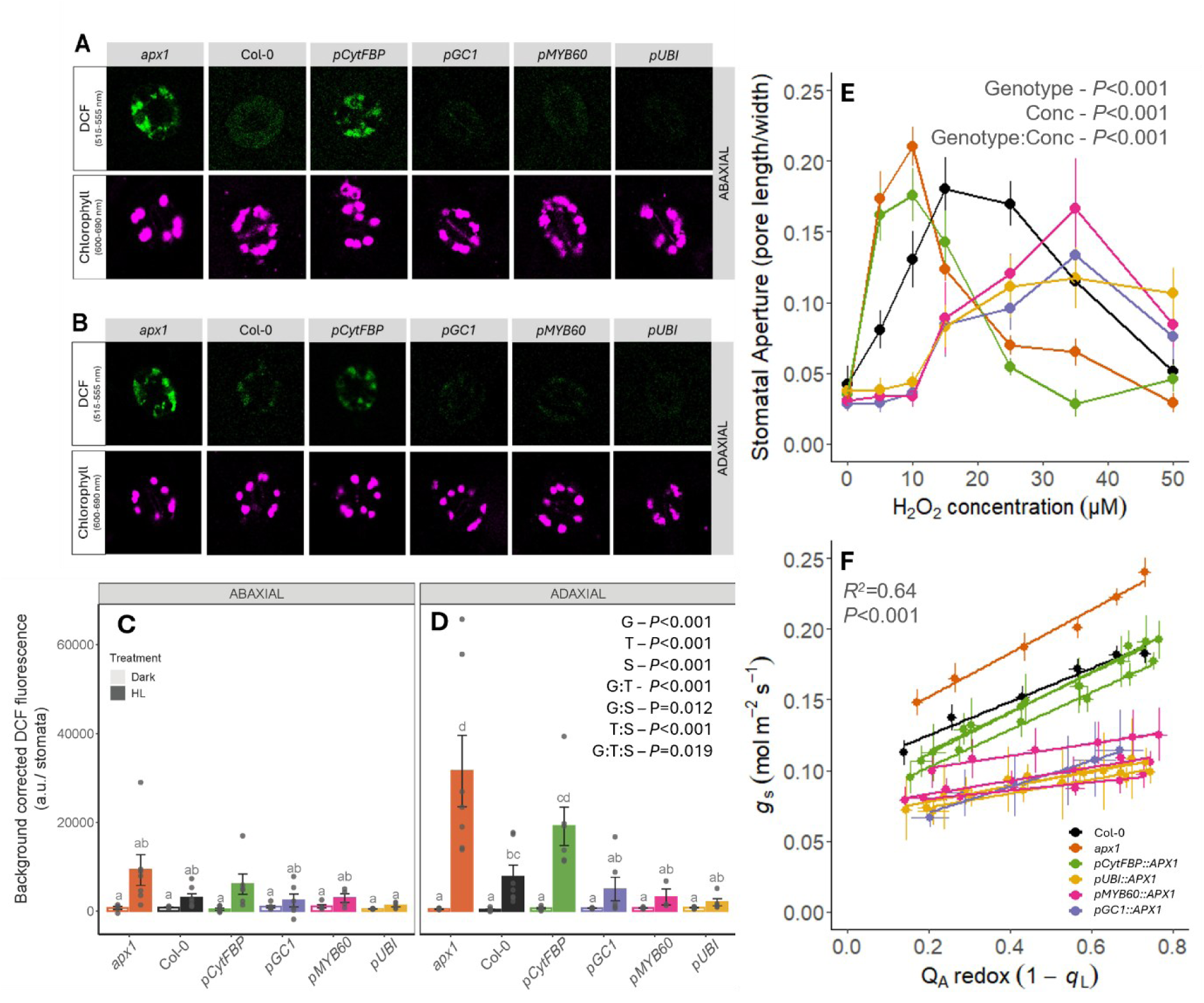
Changes in guard cell H_2_O_2_ levels and altered GC sensitivity to H_2_O_2_ in cell-specific APX1-complemented lines. Abaxial (A) and Adaxial (B) CLSM images of guard cell DCF fluorescence and chlorophyll autofluorescence. Quantification of DCF intensity in abaxial (C) and adaxial (D) guard cells, following a 2-hour high red light (800 µmol m^-2^ s^-1^; HL) treatment or dark control. n= 6-7 biological reps, >40 stomata per replicate. A three-way ANOVA was used to test the effects of genotype, leaf side and light treatment, and their interactions on the response of DCF fluorescence. Due to a significant interaction between genotype and treatment, a Tukey’s post hoc compared the genotype effects within each light treatment across leaf sides; different letters represent significant differences (*P*<0.05). All statistics were performed on log transformed DCF values to normalise data. (E) Guard cell responses to H_2_O_2_ concentration were measured in isolated epidermal peels in the absence of light. For each cell-specific complementation promoter (*pGC1, pMYB60, pCytFBPase* and *pUBI*) one representative line was used. n=5 biological replicates (>40 stomata measured per biological replicate), error bars = ±SEM. (F) Multiple linear regression analysis of the relationship between *Q*_A_ redox state (1-*q*_L_) and *g*_s_ shows that elevated guard cell H_2_O_2_ scavenging disrupts the wild type relationship. n=7, error bars = ±SEM. *Abbreviations for statistics: G = genotype; T = treatment, S = leaf side*.

To investigate the latter, guard cell sensitivity to H_2_O_2_ was assessed using dose-responses of stomatal aperture to H_2_O_2_ of epidermal peels from a representative line from each construct. As previously observed (Fig. 1K-L), stomatal aperture displayed a clear bell-shaped response to increasing H_2_O_2_ concentration (Fig. 4E), consistent with the dual effect of H_2_O_2_ on stomatal movements (i.e. stomatal opening and closing^33^). Notably, stomatal aperture was more sensitive to H_2_O_2_ concentration in *apx1* and the mesophyll-specific *apx1/pCytFBPase::APX1* lines, such that both stomatal opening and closing were shifted to lower H_2_O_2_ concentrations in these genotypes compared to Col-0. By contrast, complementation of APX1 in guard cells, either via ubiquitous or guard cell-specific promoters, showed a shift in stomatal aperture responses in the opposite direction, with strongly decreased stomatal sensitivity to low levels of H_2_O_2_,_­_ effectively shifting the concentration range of H_2_O_2_ which elicits stomatal responses upwards. The intermediate H_2_O_2_ sensitivity displayed in Col-0, shows that guard cell H_2_O_2_ scavenging capacity in pGC1, pMYB60 and pUBI lines was in excess of wild-type levels.

Finally, we evaluated the empirical correlation between Q_A_ redox state and *g*_s_ across all cell-specific *APX1*-complementation lines, which should be strongly perturbed if the link between PQ redox state changes and stomatal red light responses involves H_2_O_2_ as a signalling molecule. As expected, cell-specific complementation of *APX1* did not significantly affect values of Q_A_ redox state in response to red light (see Supplemental Fig. 6). However, previously observed correlations between Q_A_ redox state and *g_s_* (Fig. 1F) were indeed strongly affected (Fig. 4F). Multiple linear regression yielded a model for the prediction of *g*_s_ with significant main effects of *Q*_A_ redox, genotype and their interaction (*F*(23, 543)=40.65, *P*<0.001, *R*^2^=0.64; see Supplementary Table 18). While the regression intercept was significantly increased in *apx1* (*P*=0.048), both the regression intercepts and slopes were statistically similar to wild-type in all mesophyll-complemented *pCytFBPase::APX1* lines, suggesting that the mesophyll contribution to guard cell H_2_O_2_ levels in *apx1* is suppressed by mesophyll-specific APX1 complementation. The significantly decreased intercepts and regression slopes across all ubiquitous and guard cell APX1-complemented lines are consistent with strong suppression of guard cell H_2_O_2_ levels, resulting in complete disruption of the coordination between PQ redox and stomatal opening (see Supplementary Table 18 for full regression analysis).

## Discussion

Although the stomatal response to red light is often considered ‘photosynthesis-dependent’^44^, the precise mechanisms that coordinate stomatal movements with photosynthesis remain poorly understood. While early studies proposed that red light-driven stomatal movements are only indirectly coordinated with photosynthesis via changes in intercellular CO_2_ concentration (*C*­_I_)^45,46^, *g*_s_ still responds to changes in red light intensity when *C*_i_ is held constant^1,2,47^ and the amplitude of *C*_i_-independent stomatal opening is of equal magnitude to CO_2_ driven movements^3^. Here we demonstrate that H_2_O_2_ signals, an inevitable by-product of photosynthesis^10,11^, underly these responses.

In the context of stomatal regulation, most work has focused on the role of H_2_O_2_ in stomatal closure^38,48^, but our results strongly show its involvement in stomatal opening, in line with previous work^9,33^. While photosynthesis seems to provide the major fraction of H_2_O_2_ accumulation in guard cells, strong suppression of stomatal opening via the inhibition of RBOH activity, either chemically using the NADPH-oxidase inhibitors DPI and imidazole (Fig. 2A-G), or genetically in *rbohd rbohf* double mutants^9,33^, shows that RBOH-dependent generation of apoplastic H_2_O_2_ also contributes to H_2_O_2_ accumulation in response to red light, potentially as signal amplification via positive feedback^49^.

Our finding that photosynthetic H_2_O_2_ signals underpin the stomatal red light response also provides a mechanistic explanation to reconcile contradicting observations regarding the involvement of guard cells and mesophyll cells (reviewed by Lawson et al., 2014; Lawson & Matthews, 2020). H_2_O_2_ is well-known to play a role in cell-cell communication^32^ and there are multiple, non-exclusive routes by which mesophyll-derived H_2_O_2_ could be transferred to guard cells, including RBOH-dependent apoplastic H_2_O_2_ signalling, symplastic communication through plasmodesmata^29^ and diffusion through aquaporins^51^. Indeed, our results clearly demonstrate a significant effect of both mesophyll and guard cell-specific complementation of *APX1* in the *apx1* background on guard cell H_2_O_2_ accumulation (Fig. 4A-B) and stomatal opening (Fig. 3D). These findings seem consistent with involvement of both mesophyll and guard cell photosynthesis on phosphorylation of the penultimate Threonine in guard cells PM H^+^ATPases^5,52^, and possibly also Threonine-881^8^. However, the molecular nature of signals which mediate photosynthesis-dependent PM H^+^-ATPase activation still remain to be determined and could be independent from H_2_O_2_.

Finally, we demonstrate that H_2_O_2_ signalling provides the mechanistic link underpinning empirical correlations between PQ redox state changes and stomatal opening^3,36,53–55^ which can explain the positive impact of *PsbS* overexpression on intrinsic water use efficiency^36,37^. Disruption of the relationship between PQ redox state and *g*_s_ in cell-specific *apx1*/*APX1* lines (Fig. 4F) shows that this correlation is indirect, consistent with slight genotype-specific deviations of slope and intercept in rice lines with altered *PsbS* expression^55^, but since the effect of *PsbS* on H_2_O_2_ accumulation is upstream from the action of *APX1*, manipulation of scavenging capacity via *APX1* provides a more direct regulator of stomatal aperture. Perhaps as a result, the strong suppression of red light-induced stomatal opening observed in APX1 complemented lines (Fig. 3D), is substantially more pronounced than in *PsbS*-overexpressing tobacco ^36,37^ and rice^55^. Intriguingly, despite only minor suppression of red light-induced stomatal opening, *PsbS-*overexpression led to water use efficiency (WUE) improvements of 25% in tobacco^36,37^ and 11% in rice^55^. Plant WUE is a major constraint to global agriculture^56^, leading to an unsustainable demand for freshwater supplies, which is further exacerbated by climate change^57^. Genetic engineering strategies to enhance WUE in crop species are therefore urgently needed. Based on our results, manipulation of guard cell H_2_O_2_ scavenging capacity can be used to target H_2_O_2_ signalling and thereby adjust coordination between photosynthesis and stomatal conductance, which may offer potential for new WUE improvement strategies.

## Methods

### Plant Materials and Growth Conditions

Seeds of the *Arabidopsis thaliana* ascorbate peroxidase1 knock out mutants, *apx1* (*apx1-1*: SALK_000249^58^ and *apx1-2*: SALK_088569) were obtained from the Nottingham Arabidopsis Stock Centre (NASC) and genotyped via PCR (see Supplementary Table 1 for primer sequences). The *Arabidopsis Photosystem II Subunit S* overexpression line L17^35^, was donated by Professor Krishna Niyogi. Homozygous *apx1L17* double mutants were generated by crossing with *apx1-1* and genotyped via PCR. For all experiments Col-0 was used as wild-type control.

Seeds were sown on a 4:1 mix of Levington® Advance F2 compost: sand and stratified at 4°C for 4 days. Seedlings were then transferred into individual 7×7 cm pots and positioned in a controlled growth chamber with a short-day photoperiod (8 h light/16 h dark). Light intensity was controlled at ∼150 µmol m^-2^ s^-1^, relative humidity at 60% and air temperature at 20°C. Plants were hand watered and randomly repositioned on the growth shelf every 3-4 days. All experiments were performed on young fully expanded leaves of 6-8 week old plants.

### Generation of APX1-complementation lines

Cell-specific *APX1* expression constructs were generated using GoldenGate cloning^59^. Full length sequences of the guard cell specific *GC1* promoter (1700 bp; AT1G22690^40^) and *MYB60* promoter (1500 bp; AT5G05930^41^), along with the full length *AtAPX1* coding sequence (AT1G07890), were PCR amplified and domesticated via site directed mutagenesis from *Arabidopsis* DNA (see Supplementary Table 1 for primer sequences). The mesophyll-specific cytosolic fructose 1,6-bisphosphatase (*CytFBPase*) promoter and tobacco mosaic virus (TMV) 5’UTR^42,43^ were kindly provided by Professor Uwe Sonnewald (Friedrich-Alexander-University, Germany). Additional constitutive expression constructs were generated using the ubiquitin promoter (*UBI*). All constructs used the full *APX1* coding sequence and HEAT SHOCK PROTEIN 18.2 terminator (*hspT*). The *pCytFBPase*/*pGC1*/*pMYB60*/*pUBI::APX1:hspT* constructs were generated using a two-step GoldenGate reaction, first cloning into EC47742\pL1V-F2 level and subsequently into EC50505/pL2V-1 along with the FAST seed fluorescent marker (*pOLE::OLE-mRuby:35sT*) in EC47802\pL1V-R1, and transformed into *apx1-2*. To verify promoter specificity, eGFP-based promoter-reporter constructs were generated for the three cell-specific promoters. Level 2 constructs were electroporated into *Agrobacterium tumefaciens* GV3101 and transformed into the *apx1-2* (SALK_088696) genetic background using the floral dip method ^60,61^. Positive transformants were selected based on seed fluorescence and single insertion homozygous T_1_ individuals were selected based on segregation of the mRuby fluorescence signal.

### Molecular characterisation of genetic mutants

To genetically characterise *Arabidopsis* mutants, protein and mRNA were extracted from the same leaf sample (NucleoSpin RNA/Protein kit, REF740933, Macherey-Nagel GmbH & Co., Düren, Germany). For each sample, 500 ng of mRNA was reverse transcribed into cDNA using the High-Capacity cDNA Reverse Transcription kit (4368814, Thermo Fisher Scientific, Waltham, MA, USA). Quantitative reverse transcription PCR (rt-qPCR) was used to quantify *AtPsbS* and *AtAPX1* transcripts relative to the reference genes *AtActin* and *AtUbiquitin*. For a list of primers used for rt-qPCR, please refer to supplementary Table 1.

Total protein concentration was quantified using a protein quantification assay based on Karlsson *et al.,* (1994) (ref. 40967.50, Macherey-Nagel GmbH & Co., Düren, Germany). Samples containing 5 µg total protein were separated by gel electrophoresis in a 12% sodium dodecyl sulfate-polyacrylamide gel, blotted onto a 0.2 µm PVDF membrane and immuno-labelled with primary antibodies raised against either *At*PsbS (1:2000 dilution; AS09533, Agrisera, Vannas, Swedent) or *At*APX1 (1:15000 dilution; PA5-98316, Thermo Fisher Scientific, Waltham, MA, USA), followed by application with the secondary antibody, anti-RabbitHRP (1:12500 dilution, W401B, Promega, Madison, WI, USA). Chemiluminescence was detected using Clarity™ Western ECL Substrate (1705060, Bio-Rad). A PageRuler Plus Protein ladder was used as a size indicator on each gel. The blots were subsequently stained with Coomassie Brilliant Blue to check for equal protein loading.

### Gas exchange measurements

Col-0 and mutant plants were dark adapted for 2 hours, after which a fully expanded leaf (rosette leaf number 7-9) was clamped into the cuvette of an open gas exchange system (LI6400XT, LI-COR) with a 2 cm^2^ integrated fluorometer head (Leaf Chamber Fluorometer, LI6400-40, LI-COR). Block temperature was controlled at 25° C, reference CO_2_ at 410 ppm and leaf to air VPD was maintained at approximately 1.2 kPa. For leaves that did not completely fill the cuvette, an image of the leaf inside the gasket was taken and leaf area was calculated using ImageJ (ImageJ, U. S. National Institutes of Health, MD, USA) and adjusted in the gas exchange calculations. The reference CO­_2_ was adjusted to maintain a constant intercellular CO_2_ concentration (*C*_i_) of 300 µmol mol^-1^, which is the approximate steady-state *C*_i_ value measured at high red light (800 µmol m^-2^ s^-1^) for WT *Arabidopsis*^3^ Click or tap here to enter text. Leaves were allowed to equilibrate for at least 30 minutes in the dark at this *C*_i_ to ensure *g_s_* had stabilised. The measuring light was then switched on briefly and once *F* stabilised, chlorophyll fluorescence and gas exchange parameters were recorded. All fluorescence measurements were obtained using the multiphase flash routine^62^, with the saturating flash set to 4000 µmol m^-1^ s^-2^ and ramp rate of 30%. Minimal (F_o_) and maximal (F_m_) dark-adapted fluorescence yields were used to determine the maximal efficiency of whole chain electron transport (F_v_/F_m_). The actinic light (100% red light, 635 ± 10 nm full width half maximum) was then increased stepwise: 0, 50, 100, 200, 400, 600, 800 µmol m^-2^ s^-1^. For each increase in red light, the CO_2_ concentration of the cuvette was altered via the millivolt signal to maintain a constant *C*_i_ concentration of ∼300 µmol mol^-1^ across the entire red light response curve. Plants were acclimated for at least 30 minutes at each light intensity before recording gas exchange and chlorophyll fluorescence parameters (*F*’, *F*_o_’ and *F*_m_’). For the definitions and equations of the fluorescence parameters used, please refer to Supplemental Table 2.

### Measurement of H_2_O_2_ in guard cells by H_2_DCFDA

Accumulation of H_2_O_2_ in the guard cells was assayed using 2′,7′-dichlorodihydrofluorescein diacetate (H_2_DCFDA). 10 µM H_2_DCFDA was infiltrated into detached *Arabidopsis* leaves as described previously ^28^, with some modifications. In short, *Arabidopsis* leaves were petiole infiltrated with H_2_DCFDA under weak red light (20 µmol m^-2^ s^-1^) for up to 12 hours to ensure complete absorption throughout the leaf. The following morning, leaves were illuminated with high red light (HL) (800 µmol m^-2^ s^-1^, 635 nm ± 10 FWHM) for 1 hour. As a control for each HL treated replicate, a second leaf was kept in darkness (D). For both treatments, leaves were placed on damp tissue paper and sandwiched between 2 sheets of clear Perspex to prevent leaf drying. After the 1-hour treatment, abaxial and adaxial epidermal sections were isolated using the tape-peel method ^63^, washed to remove any remaining mesophyll debris, mounted onto a slide and imaged immediately via confocal laser scanning microscopy (Leica TCS SP8, Leica, Germany). DCF fluorescence was excited with the 488 nm laser at 15% power and detected at 522 (±10) nm, chlorophyll autofluorescence was detected at 675 (±25) nm. Gain was set at 300 % and 150 % for DCF and chlorophyll autofluorescence, respectively. Brightfield images were captured in parallel to identify guard cell outlines. For each biological replicate, four randomly selected images were taken per leaf side, two on either side of the midvein (image size = 291 x 291 µm).

ImageJ software (ImageJ, U. S. National Institutes of Health, MD, USA) was used to quantify H_2_O_2_ accumulation via the DCF signal. The brightfield channel was used to identify each stoma as a region of interest (ROI). The same ROIs were then applied to the chlorophyll and DCF channels, and the integrated density (mean fluorescence intensity x ROI area) of each ROI was recorded for each channel. To correct fluorescence values for background noise, an additional ROI (a non-stomatal epidermal region) was selected in each image. For each image, the DCF and chlorophyll fluorescence was background-corrected using the following equation: background corrected fluorescence = Integrated Density_stomata_ - (Mean_background_ X Area_stomata_), where Integrated Density_stomata_ is the raw fluorescence intensity of the stomatal ROI, Mean_background_ denotes the mean fluorescence intensity of the background ROI, and Area_stomata_ is the area of the stomatal ROI. For each biological replicate (n=6), mean fluorescence intensity was calculated by averaging the values obtained from the four images, encompassing >40 stomata per biological replicate for each leaf side.

### Measurement of H_2_O_2_ in whole leaves by Amplex Red

H_2_O_2_ accumulation in whole leaves was quantified using the Amplex™ Red Hydrogen Peroxide Microplate Assay Kit (Invitrogen™, A22188). Fully expanded leaves from 6- to 8-week-old *Arabidopsis thaliana* plants were detached at the petiole, placed on damp tissue paper between two sheets of clear Perspex, and dark-adapted for 2 hours. Leaves were then exposed to high red light (800 µmol m⁻² s⁻¹, 635 nm ± 10 FWHM) or kept in darkness for 1 hour. Following treatment, 6 mm leaf discs were excised, snap-frozen in liquid nitrogen, and pooled in pairs per biological replicate. Leaf tissue was ground to a fine powder using a tissue lyser, and 200 µl of Amplex™ Red reaction buffer (0.25 M sodium phosphate, pH 7.4) was added. H_2_O_2_ was extracted by shaking the samples at 25°C for 15 minutes, followed by centrifugation at 14,000 g for 10 minutes at 4°C. The supernatant was transferred to a fresh tube, centrifuged again for 2 minutes, and stored on ice. H_2_O_2_ levels were quantified in black clear bottomed 96-well plates (Invitrogen^TM^, M33089) following the manufacturer’s instructions. 10 µl of supernatant was mixed with 40 µl of reaction buffer, and the reaction was initiated by adding 50 µl of Amplex™ Red reaction mix. Samples were briefly shaken and incubated in darkness for 30 minutes, after which resorufin fluorescence was measured using a CLARIOstar plate reader (excitation: 570 nm; emission: 585 nm).

### Stomatal response to H_2_O_2_

All stomatal aperture assays were carried out using fully expanded *Arabidopsis* leaves of 6–8-week-old plants. Stomatal responses to H_2_O_2_ concentration were investigated following the protocol by Li *et al.,* (2020), with some modifications. In short, abaxial and adaxial epidermal peels were separated using the tape-peel method^63^ and pre-incubated in the stomatal assay buffer (10 mM KCl, 7.5 mM iminodiacetic acid, and 10 mM MES) for 2.5 hours in the dark. Following dark adaptation, H_2_O_2_ (30% W/V freshly diluted to 100 mM; HYD005, Thermo Fisher Scientific, Waltham, MA, USA), was added to the stomatal assay buffer to achieve the following concentrations: 0, 5, 10, 15, 25, 35, 50 µM. Epidermal peels were incubated with H_2_O_2_ for a further 2.5 hours in dark, after which samples were mounted onto a slide and imaged immediately.

### Stomatal red light response in the presence of inhibitors

Stomatal aperture responses to red light were measured in the presence of either H_2_O_2_ scavengers or inhibitors of photosynthetic electron transport. *Arabidopsis* leaves were excised at the petiole and incubated for 2 hrs in darkness in buffer containing either the H_2_O_2_ scavengers 1 mM potassium iodide (KI) or 100 µM dimethylthiourea (DMTU); the NADPH oxidase inhibitors 20 µM diphenyleneiodonium chloride (DPI) or 20 mM imidazole; the photosynthetic electron transport inhibitors 10 µM DCMU or 20 µM DBMIB; or mock solution (0.05% DMSO). After 2 hrs incubation, 20 µM H_2_DCFDA was added to the buffer and leaves were further incubated for 1 hour. Subsequently, leaves were briefly dried and treated for 1 hr with either red light (800 µmol m^-2^ s^-1^, 635 nm ± 10 FWHM) or darkness. Epidermal layers were separated as above and imaged immediately. All images were obtained within the photoperiod, between the hours of 12:30 and 16:30. Stomatal aperture was quantified using Image J. For each biological replicate, the mean stomatal aperture (stomatal pore width / stomatal pore length) for each leaf side was calculated from a total of 4 images (388 µm^2^), 2 either side of the primary vein. A total of 5 biological replicates were imaged per genotype or treatment; and for each biological replicate stomatal aperture was quantified from ∼40 stomata for each leaf side.

### Statistical analyses

For gas-exchange measurements, two-way repeated measures ANOVA was used to assess the significance of effects of red light intensity, genotype and their interaction on the response of *g*_s_ and *A*_net_, as well as the chlorophyll fluorescence parameters *F*_v_/*F_m_*, ΦPSII, NPQ and 1-*q*_L_. For models which yielded a significant interaction, a Dunnett’s test was used to compare single and double mutants to Col-0 at each light level. The same analysis was performed for the APX1-complementation lines, except a nested model was used, with the effects of line nested within genotype, followed by a Tukey’s post hoc test. For both datasets, the relationship between 1-*q*_L_ and *g*_s_ was analysed using multiple linear regression and the minimum model was obtained via backward stepwise elimination. The effect of H_2_O_2_ removal on the stomatal aperture was analysed using a one-way ANOVA. For the quantification of guard cell H_2_O_2_ levels, a three-way ANOVA was used to assess the significance of light treatment, genotype, leaf side and their interactions on the fluorescence intensity of DCF, chlorophyll autofluorescence and relative chlorophyll coverage, followed by a Tukey’s multiple comparison test. Due to a non-normal distribution, analysis of guard cell H_2_O_2_ levels was conducted on log transformed data. Likewise, for the H_2_O_2_ dose-response assay a three-way ANOVA was used to assess the significance of H_2_O_2_ concentration, genotype, leaf side and their interactions on stomatal aperture, which was followed by a Tukey’s multiple comparison test. For each ANOVA, assumptions of normality, homogeneity of variance and sphericity were evaluated using Shapiro-Wilks, Levene’s and Mauchly’s tests, respectively. All data analysis and plot generation were carried out using R 4.1.1 (R Core Team, 2021) on RStudio (Posit Team, 2022).

## Acknowledgements

This work was funded via a School of Biological Sciences doctoral training award to GT and a Gatsby foundation startup award to JK. The authors thank Ms Pauline Dussouchaut for assistance with inhibitor pilot studies, Ms Caitlin Cutts and Ms Tara Garlick for their assistance with H_2_O_2_ dose response pilot studies, and Profs Julian Hibberd, Alex Webb, Julia Davies and Tracy Lawson for helpful feedback on earlier versions of the manuscript.

## Author contributions

GT and JK conceived the work, GT performed all experiments, GT performed data analysis and interpretation with help from JW and JK, JW and GT mapped the L17 locus, GT drafted the manuscript, JK and JW revised the draft manuscript, all authors approve the submitted version.

## Data availability statement

The authors confirm that all relevant data are included in the paper and its supplementary information files. Raw image files used to estimate stomatal apertures for crosses and inhibitor experiments can be downloaded from https://doi.org/10.17863/CAM.116973 and for APX1 complemented lines https://doi.org/10.17863/CAM.116859.

## References

1. Messinger, S. M., Buckley, T. N. & Mott, K. A. Evidence for involvement of photosynthetic processes in the stomatal response to CO2. Plant Physiol 140, (2006).

2. Lawson, T., Lefebvre, S., Baker, N. R., Morison, J. I. L. & Raines, C. A. Reductions in mesophyll and guard cell photosynthesis impact on the control of stomatal responses to light and CO2. J Exp Bot 59, (2008).

3. Taylor, G., Walter, J. & Kromdijk, J. Illuminating stomatal responses to red light: establishing the role of C i-dependent versus -independent mechanisms in control of stomatal behaviour . J Exp Bot (2024) doi:10.1093/jxb/erae093.

4. Sharkey, T. D. & Raschke, K. Effect of Light Quality on Stomatal Opening in Leaves of Xanthium strumarium L. . Plant Physiol 68, (1981).

5. Ando, E. & Kinoshita, T. Red light-induced phosphorylation of plasma membrane H+-ATPase in stomatal guard cells. Plant Physiol 178, (2018).

6. Wang, S. W. et al. Lacking chloroplasts in guard cells of crumpled leaf attenuates stomatal opening: Both guard cell chloroplasts and mesophyll contribute to guard cell ATP levels. Plant Cell Environ 37, (2014).

7. Fuji, S. et al. Light-induced stomatal opening requires phosphorylation of the C-terminal autoinhibitory domain of plasma membrane H+-ATPase. Nat Commun 15, (2024).

8. Hayashi, Y. et al. Phosphorylation of plasma membrane H+-ATPase Thr881 participates in light-induced stomatal opening. Nat Commun 15, (2024).

9. Shi, W. et al. Hydrogen peroxide is required for light-induced stomatal opening across different plant species. Nat Commun 15, (2024).

10. Asada, K. Production and scavenging of reactive oxygen species in chloroplasts and their functions. Plant Physiology vol. 141 Preprint at 10.1104/pp.106.082040 (2006).

11. Foyer, C. H. & Hanke, G. ROS production and signalling in chloroplasts: cornerstones and evolving concepts. Plant Journal 111, (2022).

12. Perry, J. J. P., Shin, D. S., Getzoff, E. D. & Tainer, J. A. The structural biochemistry of the superoxide dismutases. Biochimica et Biophysica Acta - Proteins and Proteomics vol. 1804 Preprint at 10.1016/j.bbapap.2009.11.004 (2010).

13. Pilon, M., Ravet, K. & Tapken, W. The biogenesis and physiological function of chloroplast superoxide dismutases. Biochimica et Biophysica Acta - Bioenergetics vol. 1807 Preprint at 10.1016/j.bbabio.2010.11.002 (2011).

14. Mubarakshina, M. M. & Ivanov, B. N. The production and scavenging of reactive oxygen species in the plastoquinone pool of chloroplast thylakoid membranes. Physiologia Plantarum vol. 140 Preprint at 10.1111/j.1399-3054.2010.01391.x (2010).

15. Mubarakshina, M. M. et al. Production and diffusion of chloroplastic H2O2 and its implication to signalling. J Exp Bot 61, (2010).

16. Yadav, D. K., Prasad, A., Kruk, J. & Pospíšil, P. Evidence for the involvement of loosely bound plastosemiquinones in superoxide anion radical production in photosystem II. PLoS One 9, (2014).

17. Khorobrykh, S. A., Karonen, M. & Tyystjärvi, E. Experimental evidence suggesting that H2O2 is produced within the thylakoid membrane in a reaction between plastoquinol and singlet oxygen. FEBS Lett 589, (2015).

18. Mubarakshina, M., Khorobrykh, S. & Ivanov, B. Oxygen reduction in chloroplast thylakoids results in production of hydrogen peroxide inside the membrane. Biochim Biophys Acta Bioenerg 1757, (2006).

19. Mubarakshina, M. M., Khorobrykh, S. A., Kozuleva, M. A. & Ivanov, B. N. Intramembrane formation of hydrogen peroxide during oxygen reduction in thylakoids of higher plants. Dokl Biochem Biophys 408, (2006).

20. Smirnoff, N. & Arnaud, D. Hydrogen peroxide metabolism and functions in plants. New Phytologist vol. 221 Preprint at 10.1111/nph.15488 (2019).

21. Bienert, G. P., Schjoerring, J. K. & Jahn, T. P. Membrane transport of hydrogen peroxide. Biochimica et Biophysica Acta - Biomembranes vol. 1758 Preprint at 10.1016/j.bbamem.2006.02.015 (2006).

22. Ślesak, I., Libik, M., Karpinska, B., Karpinski, S. & Miszalski, Z. The role of hydrogen peroxide in regulation of plant metabolism and cellular signalling in response to environmental stresses. Acta Biochimica Polonica vol. 54 Preprint at 10.18388/abp.2007_3267 (2007).

23. Exposito-Rodriguez, M., Laissue, P. P., Yvon-Durocher, G., Smirnoff, N. & Mullineaux, P. M. Photosynthesis-dependent H2O2 transfer from chloroplasts to nuclei provides a high-light signalling mechanism. Nat Commun 8, (2017).

24. Lim, S. L. et al. Arabidopsis guard cell chloroplasts import cytosolic ATP for starch turnover and stomatal opening. Nat Commun 13, (2022).

25. Roelfsema, M. R. G. & Hedrich, R. In the light of stomatal opening: New insights into ‘the Watergate’. New Phytologist vol. 167 Preprint at 10.1111/j.1469-8137.2005.01460.x (2005).

26. Mott, K. A., Sibbernsen, E. D. & Shope, J. C. The role of the mesophyll in stomatal responses to light and CO2. Plant Cell Environ 31, (2008).

27. Lawson, T., Simkin, A. J., Kelly, G. & Granot, D. Mesophyll photosynthesis and guard cell metabolism impacts on stomatal behaviour. New Phytologist vol. 203 Preprint at 10.1111/nph.12945 (2014).

28. Xiong, H. et al. Photosynthesis-independent production of reactive oxygen species in the rice bundle sheath during high light is mediated by NADPH oxidase. Proc Natl Acad Sci U S A 118, (2021).

29. Fichman, Y., Myers, R. J., Grant, D. A. G. & Mittler, R. Plasmodesmata-localized proteins and ROS orchestrate light-induced rapid systemic signaling in Arabidopsis. Sci Signal 14, (2021).

30. Rodrigues, O. et al. Aquaporins facilitate hydrogen peroxide entry into guard cells to mediate ABA- and pathogen-triggered stomatal closure. Proc Natl Acad Sci U S A 114, (2017).

31. Mott, K. A., Berg, D. G., Hunt, S. M. & Peak, D. Is the signal from the mesophyll to the guard cells a vapour-phase ion? Plant Cell Environ 37, (2014).

32. Fichman, Y. & Mittler, R. Rapid systemic signaling during abiotic and biotic stresses: is the ROS wave master of all trades? Plant Journal vol. 102 Preprint at 10.1111/tpj.14685 (2020).

33. Li, J. G. et al. Brassinosteroid and Hydrogen Peroxide Interdependently Induce Stomatal Opening by Promoting Guard Cell Starch Degradation. Plant Cell 32, (2020).

34. Davletova, S. et al. Cytosolic ascorbate peroxidase 1 is a central component of the reactive oxygen gene network of Arabidopsis. Plant Cell 17, (2005).

35. Li, X. P., Müller-Moulé, P., Gilmore, A. M. & Niyogi, K. K. PsbS-dependent enhancement of feedback de-excitation protects photosystem II from photoinhibition. Proc Natl Acad Sci U S A 99, (2002).

36. Głowacka, K. et al. Photosystem II Subunit S overexpression increases the efficiency of water use in a field-grown crop. Nat Commun 9, (2018).

37. Turc, B. et al. Up-regulation of non-photochemical quenching improves water use efficiency and reduces whole-plant water consumption under drought in Nicotiana tabacum. J Exp Bot 75, (2024).

38. Rodrigues, O. & Shan, L. Stomata in a state of emergency: H2O2 is the target locked. Trends in Plant Science vol. 27 Preprint at 10.1016/j.tplants.2021.10.002 (2022).

39. Manders, E. M. M., Verbeek, F. J. & Aten, J. A. Measurement of co-localization of objects in dual-colour confocal images. J Microsc 169, (1993).

40. Yang, Y., Costa, A., Leonhardt, N., Siegel, R. S. & Schroeder, J. I. Isolation of a strong Arabidopsis guard cell promoter and its potential as a research tool. Plant Methods 4, (2008).

41. Rusconi, F. et al. The Arabidopsis thaliana MYB60 promoter provides a tool for the spatio-temporal control of gene expression in stomatal guard cells. J Exp Bot 64, (2013).

42. Ebneth, M. xpressionsanalyse des Promotors einer cytosolischen Fruktose-1, 6-bisphosphatase aus Kartoffel in transgenen Tabak-und Kartoffelpflanzen. (1996).

43. Wigger, J. et al. Prevention of stomatal closure by immunomodulation of endogenous abscisic acid and its reversion by abscisic acid treatment: Physiological behaviour and morphological features of tobacco stomata. Planta 215, (2002).

44. Matthews, J. S. A., Vialet-Chabrand, S. & Lawson, T. Role of blue and red light in stomatal dynamic behaviour. Journal of Experimental Botany vol. 71 Preprint at 10.1093/jxb/erz563 (2020).

45. Heath, O. V. S. & Milthorpe, F. L. Studies in stomatal behaviour: V. The role of carbon dioxide in the light response of stomata. J Exp Bot 1, (1950).

46. Roelfsema, M. R. G., Hanstein, S., Felle, H. H. & Hedrich, R. CO2 provides an intermediate link in the red light response of guard cells. Plant Journal 32, (2002).

47. Wang, Y., Noguchi, K. & Terashima, I. Photosynthesis-dependent and -independent responses of stomata to blue, red and green monochromatic light: Differences between the normally oriented and inverted leaves of sunflower. Plant and Cell Physiology vol. 52 Preprint at 10.1093/pcp/pcr005 (2011).

48. Sierla, M., Waszczak, C., Vahisalu, T. & Kangasjärvi, J. Reactive oxygen species in the regulation of stomatal movements. Plant Physiology vol. 171 Preprint at 10.1104/pp.16.00328 (2016).

49. Kimura, S. et al. Protein phosphorylation is a prerequisite for the Ca 2+-dependent activation of Arabidopsis NADPH oxidases and may function as a trigger for the positive feedback regulation of Ca 2+ and reactive oxygen species. Biochim Biophys Acta Mol Cell Res 1823, (2012).

50. Lawson, T. & Matthews, J. Guard Cell Metabolism and Stomatal Function. Annual Review of Plant Biology vol. 71 Preprint at 10.1146/annurev-arplant-050718-100251 (2020).

51. Bienert, G. P. & Chaumont, F. Aquaporin-facilitated transmembrane diffusion of hydrogen peroxide. Biochimica et Biophysica Acta - General Subjects vol. 1840 Preprint at 10.1016/j.bbagen.2013.09.017 (2014).

52. Ando, E., Kollist, H., Fukatsu, K., Kinoshita, T. & Terashima, I. Elevated CO2 induces rapid dephosphorylation of plasma membrane H+-ATPase in guard cells. New Phytologist 236, (2022).

53. Busch, F. A. Opinion: The red-light response of stomatal movement is sensed by the redox state of the photosynthetic electron transport chain. Photosynthesis Research vol. 119 Preprint at 10.1007/s11120-013-9805-6 (2014).

54. Kromdijk, J., Głowacka, K. & Long, S. P. Predicting light-induced stomatal movements based on the redox state of plastoquinone: theory and validation. Photosynth Res 141, (2019).

55. Patel-Tupper, D. et al. Multiplexed CRISPR-Cas9 mutagenesis of rice PSBS1 noncoding sequences for transgene-free overexpression. Sci Adv 10, (2024).

56. Leakey, A. D. B. et al. Water Use Efficiency as a Constraint and Target for Improving the Resilience and Productivity of C 3 and C 4 Crops. Annual Review of Plant Biology vol. 70 Preprint at 10.1146/annurev-arplant-042817-040305 (2019).

57. Yang, H., Sun, H., Jia, C., Yang, T. & Yang, X. Future Climatic Projections and Hydrological Responses with a Data Driven Method: A Regional Climate Model Perspective. Water Resources Management 38, (2024).

58. Suzuki, N., Miller, G., Sejima, H., Harper, J. & Mittler, R. Enhanced seed production under prolonged heat stress conditions in Arabidopsis thaliana plants deficient in cytosolic ascorbate peroxidase 2. J Exp Bot 64, (2013).

59. Engler, C., Kandzia, R. & Marillonnet, S. A one pot, one step, precision cloning method with high throughput capability. PLoS One 3, (2008).

60. Clough, S. J. & Bent, A. F. Floral dip: A simplified method for Agrobacterium-mediated transformation of Arabidopsis thaliana. Plant Journal 16, (1998).

61. Zhang, X., Henriques, R., Lin, S. S., Niu, Q. W. & Chua, N. H. Agrobacterium-mediated transformation of Arabidopsis thaliana using the floral dip method. Nat Protoc 1, (2006).

62. Loriaux, S. D. et al. Closing in on maximum yield of chlorophyll fluorescence using a single multiphase flash of sub-saturating intensity. Plant Cell Environ 36, (2013).

63. Wu, F. H. et al. Tape-arabidopsis sandwich - A simpler arabidopsis protoplast isolation method. Plant Methods 5, (2009).

