## Supplemental Figures 1-6 for "Chloroplast-derived hydrogen peroxide coordinates photosynthesis with stomatal opening"

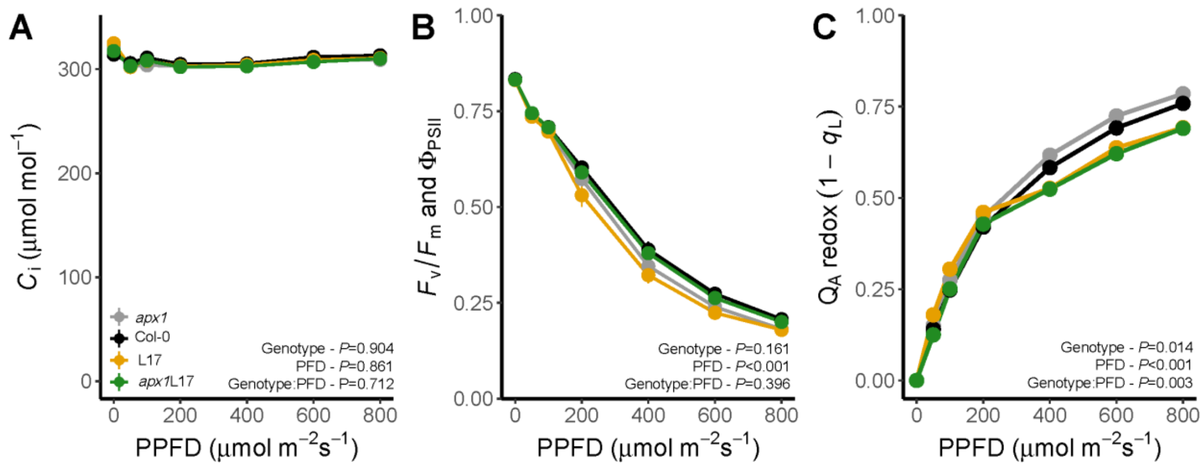


**Supplementary Figure 1. Integrated gas exchange and chlorophyll fluorescence parameters measured under constant intercellular $CO_2$ concentration ($C_i$).** (A) $C_i$ was controlled at approximately 300 µmol mol$^{-1}$, which is approximately equal to the $C_i$ value for Col-0 measured at high red light (Taylor *et al*., 2024). (B) Maximum efficiency of PSII ($F_v/F_m$) after dark adaptation and quantum efficiency of PSII ($\Phi_{PSII}$) in response to increasing red light was unaffected by PsbS or APX1 levels. (C) The redox state of quinone A ($Q_A$ redox), as estimated by the fluorescence parameter 1-$q_L$ (Kramer *et al*., 2004) separated in a PsbS dependent manner at high red light. n=7, error bars = SEM. Two-way repeated measures ANOVA tested the effects of genotype, red light intensity (PPFD) and their interaction on each parameter.

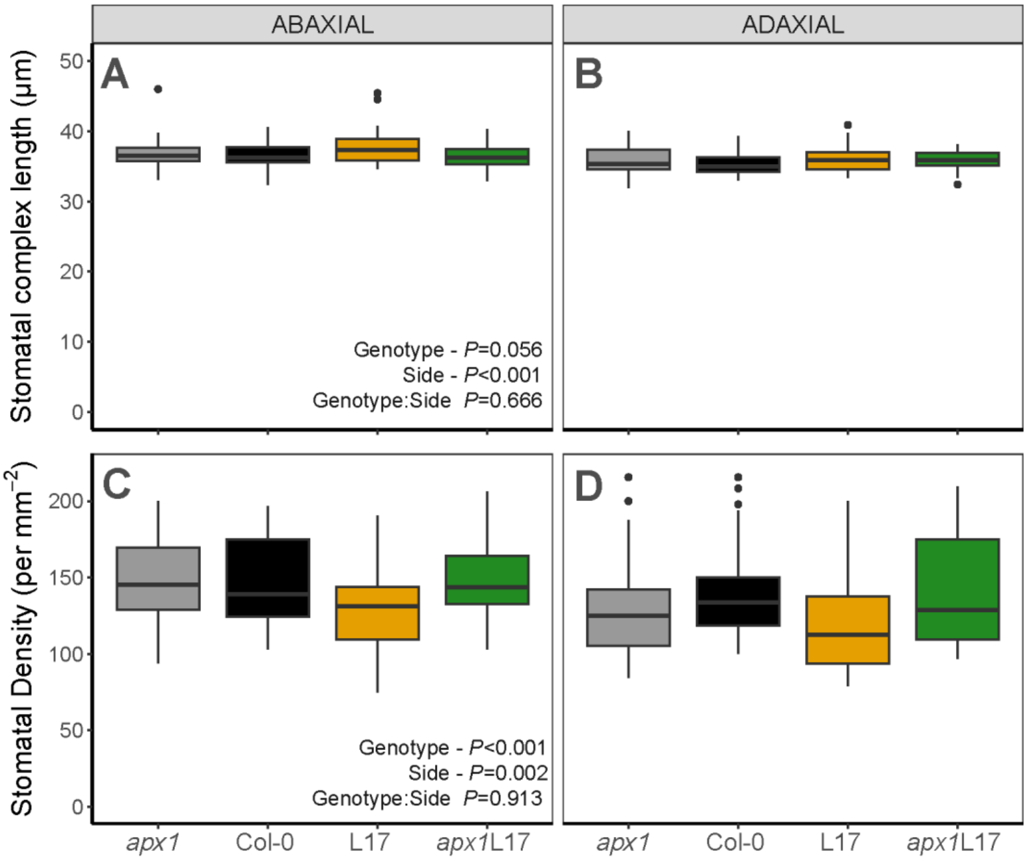


**Supplementary Figure 2. Stomatal anatomy traits do not explain variations in stomatal conductance between genotypes.** Measurements of stomatal complex length (µm) of abaxial (A) and adaxial (B) guard cells, and stomatal density (per $mm^2$) in abaxial (C) and adaxial (D) surfaces. n=10-12 independent biological replicates per genotype, >40 stomata measured per biological replicate for each leaf side. Two-way ANOVA tested the effects of genotype, leaf side and their interaction on stomatal complex length and number.

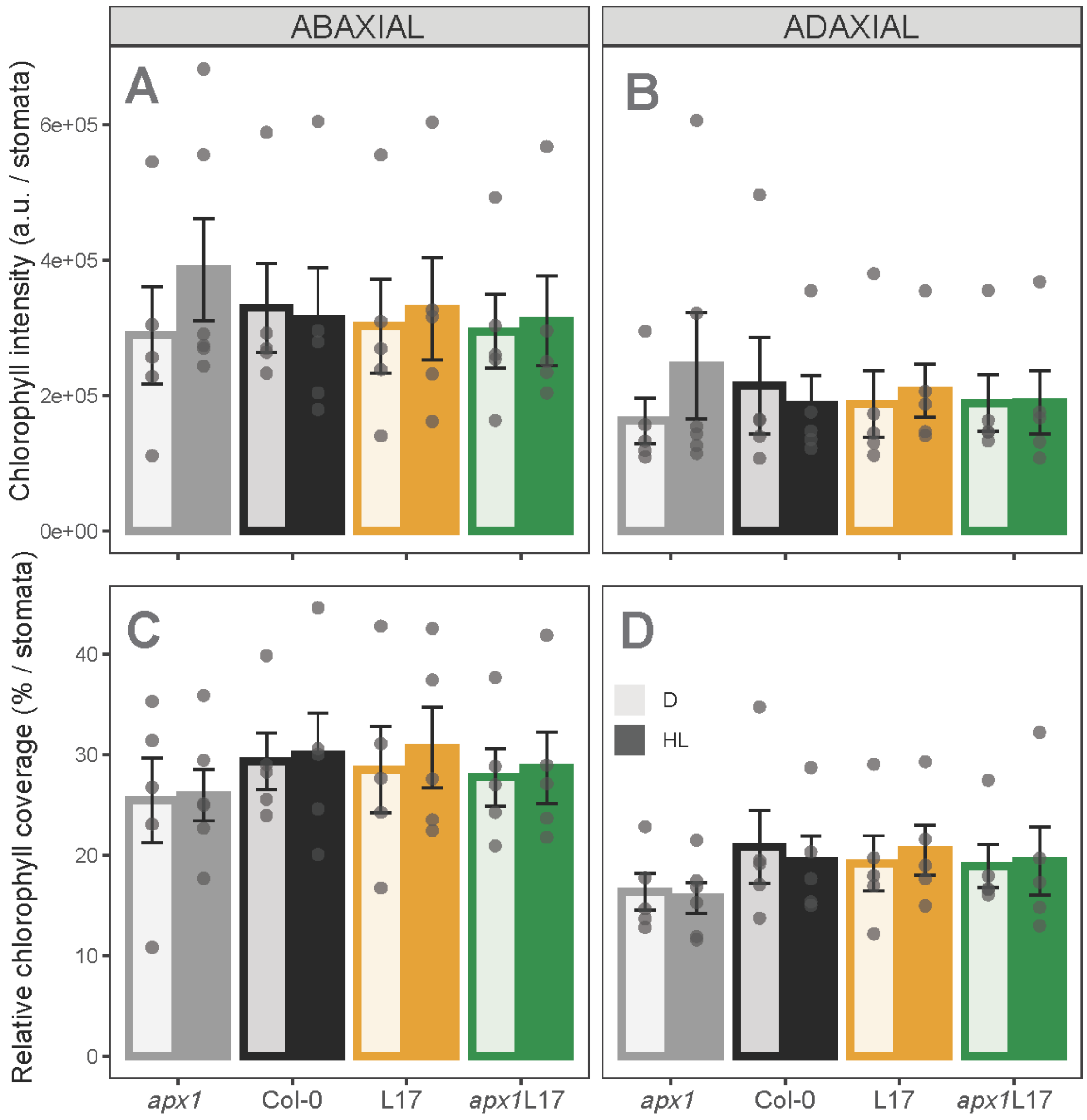


**Supplementary Figure 3. Genotypic variation in guard cell $H_2O_2$ could not be explained by differences in guard cell chloroplast coverage**. Chlorophyll fluorescence intensity (A-B) and relative chlorophyll coverage (% of stomatal area) (C-D), which provides an estimate for chloroplast area, were quantified for abaxial and adaxial guard cells from confocal laser scanning microscopy images in response to a 1-hour treatment with darkness (D) or red light illumination (HL) . Error bars represent ± SEM, n=5, >200 stomata per genotype. Three-way ANOVA tested the effects of genotype (G), light treatment (T), leaf side (S), and their interactions on guard cell chlorophyll coverage intensity and fluorescence intensity.

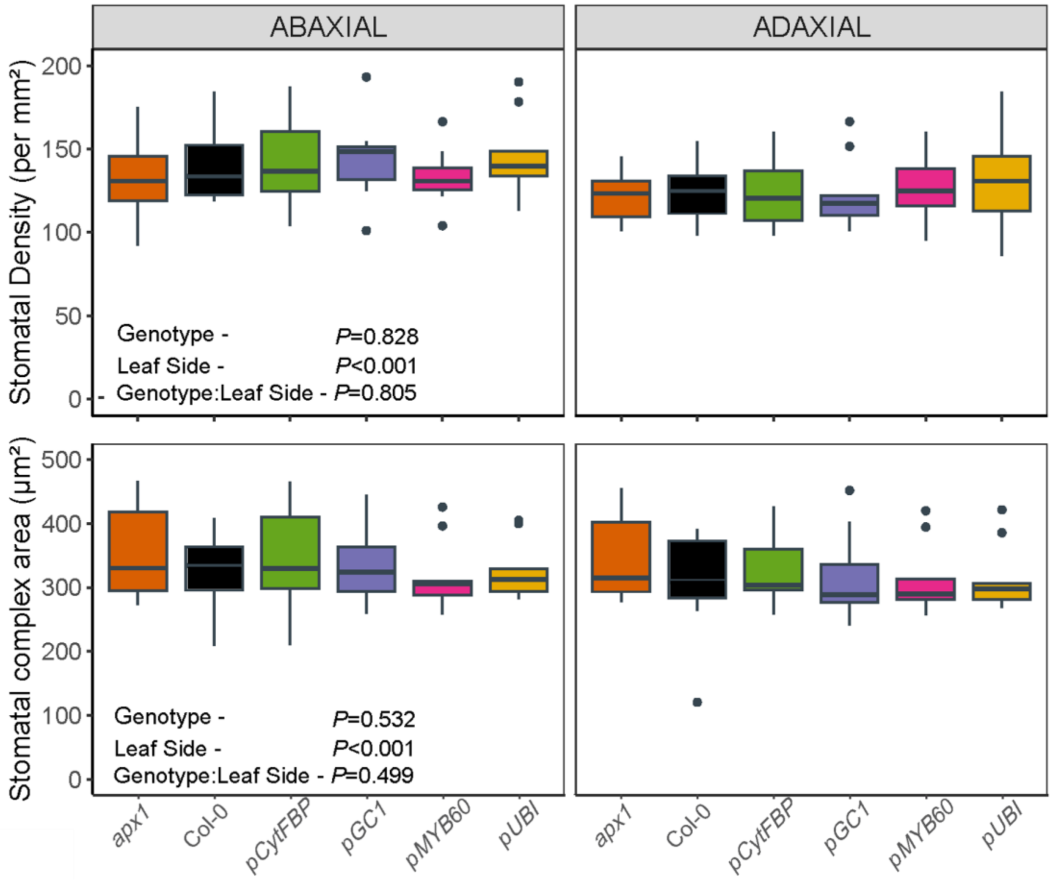


**Supplemental Figure 4. Physical stomatal characteristics of APX1-complemented plants do not explain the variation in $g_s$.** Stomatal density (A) and stomatal complex area (B) were measured in abaxial and adaxial epidermal peels. n=10-12 independent biological replicates per genotype. >40 stomata measured per biological replicate for each leaf side. Two-way ANOVA tested the effects of genotype, leaf side and their interaction on stomatal length and complex size.

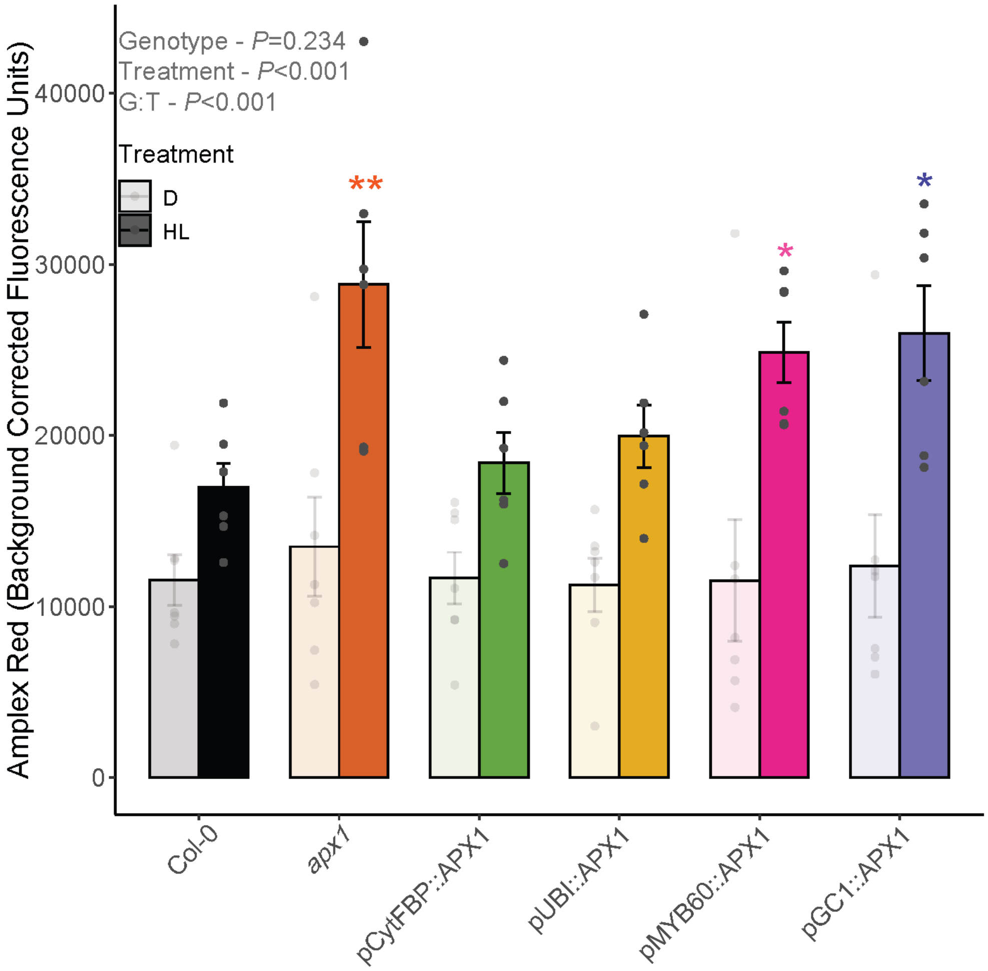


**Supplemental Figure 5. Quantification of $H_2O_2$ using Amplex Red from whole leaf discs confirms APX1 complementation in targeted cell types.** Whole leaf $H_2O_2$ concentration was quantified via Amplex Red fluorescence in response to a 1-hr red light treatment (HL; 635 nm, 800 µmol $m^{-2}$ $s^{-1}$) or darkness (D). Two-way ANOVA tested the effect of genotype, light treatment and their interaction (G:T) on the intensity of Amplex Red fluorescence. Due to a significant interaction, the effect of genotype was compared within each light treatment, asterisks represent significant differences compared to Col-0 using Dunnett's multi-comparison test (n=8, error bars represent SEM).

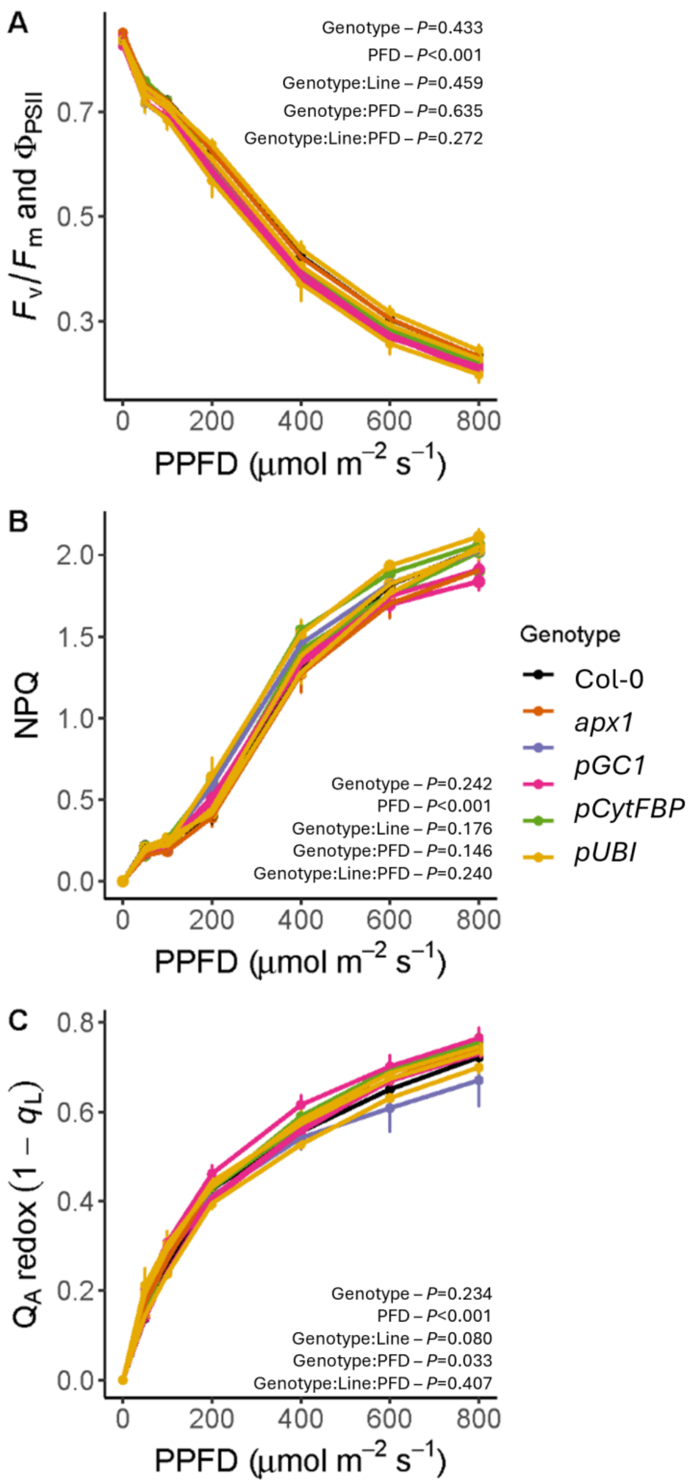


**Supplemental Figure 6. Complementation of *apx1* with cell type-specific *APX1* did not alter the response of photosynthetic electron transport to red light intensity.** The chlorophyll fluorescence parameters (A) maximum quantum yield of PSII (*Fv/Fm*) and PSII operating efficiency (ΦPSII), (B) non photochemical quenching (NPQ) and (C) the redox state of quinone A (1 – $q_L$; Kramer *et al.,* 2004) were measured in response to increasing red light intensity concomitantly alongside gas exchange. Two-way repeated measures ANOVA tested the effects of genotype, red light intensity (PPFD), and their interaction on all parameters, with nested effects of line within genotype.
